## Supporting Text for "Fine-mapping from summary data with the “Sum of Single Effects” model"

### Detailed methods

#### IBSS-ss algorithm

*SuSiE-RSS* uses the IBSS-ss algorithm, which is essentially the IBSS algorithm [1] modified for sufficient statistics or summary statistics. When applied to sufficient statistics, the IBSS-ss algorithm returns the same results as the IBSS algorithnm applied to the individual-level data. In this section we describe the IBSS-ss algorithm in detail.

##### The single effect regression (SER) model with summary statistics

The single effect regression (SER) model is defined in [1] (see also [2]) as a multiple regression model in which exactly one variable has a non-zero effect on the outcome. It is a special case of the *SuSiE* model when $L = 1$. Posterior computations with the SER model form the basis for the *SuSiE* model fitting algorithm, IBSS, hence form the basis for IBSS-ss. We show here that posterior quantities under the SER model can be computed using summary statistics $\boldsymbol{X}^\intercal \boldsymbol{X}$ and $\boldsymbol{X}^\intercal \boldsymbol{y}$.

Formally, the SER model can be written as

$$\boldsymbol{y} = \boldsymbol{X}\boldsymbol{b} + \boldsymbol{e} \tag{28}$$
$$\boldsymbol{e} \sim \mathcal{N}_N(0, \sigma^2 \boldsymbol{I}_N) \tag{29}$$
$$\boldsymbol{b} = \boldsymbol{\gamma} b \tag{30}$$
$$\boldsymbol{\gamma} \sim \text{Multinomial}(1, \boldsymbol{\pi}) \tag{31}$$
$$b \sim \mathcal{N}(0, \sigma_0^2). \tag{32}$$

Here, $\boldsymbol{y} = (y_1, \ldots, y_N)^\intercal \in \mathbb{R}^N$ denotes the phenotypes of $N$ individuals, $\boldsymbol{X} \in \mathbb{R}^{N \times J}$ denotes their corresponding genotypes at $J$ genetic variants (SNPs), $\boldsymbol{b} = (b_1, \ldots, b_J)^\intercal$ denotes a vector of regression coefficients, $\boldsymbol{e}$ is an $N$-vector of error terms, $\sigma^2 > 0$ is the residual variance parameter, and $\boldsymbol{I}_N$ is the $N \times N$ identity matrix. To simplify the presentation, we assume $\boldsymbol{y}$ and the columns of $\boldsymbol{X}$ are centered to have mean zero, which avoids the need for an intercept term [3]. $\mathcal{N}(\mu, \sigma^2)$ denotes the univariate normal distribution with mean $\mu$ and variance $\sigma^2$, $\mathcal{N}_r(\boldsymbol{\mu}, \boldsymbol{\Sigma})$ denotes the $r$-variate normal distribution with mean $\boldsymbol{\mu}$ and variance $\boldsymbol{\Sigma}$, and Multinomial$(n, \boldsymbol{p})$ denotes the multinomial distribution with $n$ trials and category probabilities $\boldsymbol{p} = (p_1, \ldots, p_J)$. Thus,

$\boldsymbol{\gamma} = (\gamma_1, \ldots, \gamma_J) \in \{0, 1\}^J$ is a binary vector of length $J$ in which exactly one element is 1 and the rest are 0, and so $\boldsymbol{b}$ is a vector with exactly one non-zero element (except for the special case when $b = 0$). The scalar $b$ represents the value of the one non-zero element in $\boldsymbol{b}$ (the "single effect"). The prior inclusion probabilities, $\boldsymbol{\pi} = (\pi_1, \ldots, \pi_J)$, which we assume are fixed and known, give the prior probability that each genetic variant has the non-zero effect. The prior variance of the single effect, $\sigma_0^2$, and the residual variance, $\sigma^2$, are hyperparameters that can be pre-specified or, more commonly, estimated.

Given settings of the hyperparameters $\sigma^2, \sigma_0^2$, the posterior distribution of $\boldsymbol{b}$ under the SER model is worked out in [1]. We summarize it here, with a focus on performing computations using $\boldsymbol{X}^\intercal \boldsymbol{X}$ and $\boldsymbol{X}^\intercal \boldsymbol{y}$.

**Proposition 1.** Consider the SER model (28–32) with known $\sigma_0^2$ and $\sigma^2$. The posterior distribution of $\boldsymbol{b} = \boldsymbol{\gamma} b$ can be expressed in terms of univariate least-squares estimates of $b_j$, $\hat{b}_j := \boldsymbol{x}_j^\intercal \boldsymbol{y} / \boldsymbol{x}_j^\intercal \boldsymbol{x}_j$, and their variances, $s_j^2 := \sigma^2 / \boldsymbol{x}_j^\intercal \boldsymbol{x}_j$. Specifically, the posterior distribution of $\boldsymbol{\gamma}$ is

$$\boldsymbol{\gamma} \mid \boldsymbol{X}, \boldsymbol{y}, \sigma^2, \sigma_0^2 \sim \text{Multinomial}(1, \boldsymbol{\alpha}) \tag{33}$$

and the posterior distribution of $b$ given $\boldsymbol{\gamma}$ is

$$b \mid \boldsymbol{X}, \boldsymbol{y}, \sigma^2, \sigma_0^2, \gamma_j = 1 \sim \mathcal{N}(\mu_{1j}, \sigma_{1j}^2), \tag{34}$$

where

$$\sigma_{1j}^2 := \frac{1}{1/\sigma_0^2 + 1/s_j^2} \tag{35}$$

$$\mu_{1j} := \sigma_{1j}^2 \hat{b}_j / s_j^2 \tag{36}$$

$$\alpha_j := \frac{\pi_j \text{BF}_j}{\sum_{j'=1}^J \pi_{j'} \text{BF}_{j'}} \tag{37}$$

$$\begin{aligned} \text{BF}_j &:= \text{BF}(\boldsymbol{x}_j, \boldsymbol{y}; \sigma^2, \sigma_0^2) \\ &:= \frac{p(\boldsymbol{y} \mid \boldsymbol{x}_j, \sigma^2, \sigma_0^2)}{p(\boldsymbol{y} \mid \boldsymbol{x}_j; \sigma^2, b = 0)} \\ &= \sqrt{\frac{s_j^2}{\sigma_0^2 + s_j^2}} \times \exp\left(\frac{\hat{b}_j^2}{2 s_j^2} \times \frac{\sigma_0^2}{\sigma_0^2 + s_j^2}\right). \end{aligned} \tag{38}$$

Note that the $\alpha_j$'s (37) are the PIPs (2) under the SER model; $\text{PIP}_j = \alpha_j$, $j = 1, \ldots, J$.

Proposition 1 shows that, given $\sigma^2$ and $\sigma_0^2$, the posterior for $\boldsymbol{b}$ under the SER model can be computed from the least-squares estimates $\hat{b}_j$ and variances $s_j^2$, which in turn can be computed from $\boldsymbol{X}^\intercal \boldsymbol{X}$ and $\boldsymbol{X}^\intercal \boldsymbol{y}$. We define SER-ss as the function that returns the posterior distribution of $\boldsymbol{b}$ under the SER model given these statistics:

$$\text{SER-ss}(\boldsymbol{X}^\intercal \boldsymbol{X}, \boldsymbol{X}^\intercal \boldsymbol{y}; \sigma^2, \sigma_0^2) := (\boldsymbol{\alpha}, \boldsymbol{\mu}_1, \boldsymbol{\sigma}_1^2), \tag{39}$$

where $\boldsymbol{\mu}_1 := (\mu_{11}, \ldots, \mu_{1J})^\intercal$, $\boldsymbol{\sigma}_1^2 := (\sigma_{11}^2, \ldots, \sigma_{1J}^2)$ and $\boldsymbol{\alpha} := (\alpha_1, \ldots, \alpha_J)$.

**Remark 1.** Although we write the posterior SER-ss as a function of $\boldsymbol{X}^\intercal \boldsymbol{X}$, it actually only depends on the diagonal elements of this matrix.

Likewise, the likelihood for $\sigma_0^2$ and $\sigma^2$ under the SER model can be computed using

**Algorithm 1** Iterative Bayesian stepwise selection using sufficient or summary statistics (IBSS-ss)

**Require:** Sufficient statistics $\boldsymbol{X}^\intercal\boldsymbol{X}, \boldsymbol{X}^\intercal\boldsymbol{y}$ and, optionally, $\boldsymbol{y}^\intercal\boldsymbol{y}$, $N$ (for estimating $\sigma^2$). Alternatively, require summary data that can be used to recover these sufficient statistics, either exactly or approximately. For example, if $\boldsymbol{X}, \boldsymbol{y}$ are standardized, $\boldsymbol{X}^\intercal\boldsymbol{X}, \boldsymbol{X}^\intercal\boldsymbol{y}$ can be recovered from $\hat{\boldsymbol{R}}, \hat{\boldsymbol{z}}, N$.
**Require:** Number of effects, $L$; initial estimates of hyperparameters $\sigma^2, \boldsymbol{\sigma}_0^2$.
**Require:** Initial estimates of the posterior mean single effects, $\bar{\boldsymbol{b}}_l$, for $l = 1, \ldots, L$.

```
repeat
    ρ̄ ← Xᵀy − XᵀX Σ_{l=1}^L b̄_l                        ▷ Compute expected residuals.
    for l in 1, ..., L do
        ρ̄_l ← ρ̄ + XᵀX b̄_l                               ▷ Disregard lth single effect in residuals.
        σ²_0l ← argmax_{σ²_0} ℓ_SER-ss(σ², σ²_0; XᵀX, ρ̄)   ▷ Update σ²_0l (optional).
        (α_l, μ_1l, σ²_1l) ← SER-ss(XᵀX, ρ̄; σ², σ²_0l)      ▷ Fit SER to residuals.
        b̄_l ← α_l ∘ μ_1l                                  ▷ "∘" denotes element-wise multiplication.
        b̄²_l ← α_l ∘ (μ_1l ∘ μ_1l + σ²_1l)                 ▷ Compute posterior second moments.
        ρ̄ ← ρ̄_l − XᵀX b̄_l                                ▷ Update expected residuals.
    end for
    σ² ← (1/N) ERSS-ss(XᵀX, Xᵀy, yᵀy, b̄_1, ..., b̄_l, b̄²_1, ..., b̄²_L)   ▷ Optional.
until convergence criterion is satisfied
return α_1, μ_11, σ²_11, ..., α_L, μ_1L, σ²_1L.
```

only the sufficient statistics since it can be expressed as a weighted sum of the BFs:

$$\begin{aligned}\ell_{\mathrm{SER}}(\sigma^2, \sigma_0^2; \boldsymbol{X}, \boldsymbol{y}) &:= p(\boldsymbol{y} \mid \boldsymbol{X}, \sigma^2, \sigma_0^2) \\ &= p(\boldsymbol{y} \mid \boldsymbol{X}, \sigma^2, b = 0) \textstyle\sum_{j=1}^J \pi_j \mathrm{BF}_j \\ &= \mathcal{N}_N(\boldsymbol{y}; \boldsymbol{0}, \sigma^2 \boldsymbol{I}_N) \textstyle\sum_{j=1}^J \pi_j \mathrm{BF}_j \\ &:= \ell_{\text{SER-ss}}(\sigma^2, \sigma_0^2; \boldsymbol{X}^\intercal\boldsymbol{X}, \boldsymbol{X}^\intercal\boldsymbol{y}). \end{aligned} \tag{40}$$

Following [1], we compute the maximum-likelihood estimate of $\sigma_0^2$ by maximizing this likelihood via numerical optimization.

##### The IBSS-ss algorithm

The IBSS-ss algorithm is given in Algorithm 1. Additional notation used in Algorithm 1 includes: $\bar{\boldsymbol{b}}_l$, the expected value of $\boldsymbol{b}_l$ with respect to the approximate posterior distribution, $q(\boldsymbol{b})$; and $\overline{\boldsymbol{b}_l^2} = (\overline{b_{l1}^2}, \ldots, \overline{b_{lJ}^2})^\intercal$, the vector of posterior second moments $\overline{b_{lj}^2} := \mathbb{E}_q[b_{lj}^2]$. A key change in implementation is that the original IBSS algorithm keeps track of the posterior mean residuals $\bar{\boldsymbol{r}} := \mathbb{E}_q[\boldsymbol{X}^\intercal\boldsymbol{y} - \boldsymbol{b}] = \boldsymbol{X}^\intercal\boldsymbol{y} - \sum_{l=1}^L \bar{\boldsymbol{b}}_l$ whereas IBSS-ss updates $\bar{\boldsymbol{\rho}} := \boldsymbol{X}^\intercal\bar{\boldsymbol{r}}$. See [1] for development and justification for the IBSS algorithm, and details of implementation, including preprocessing steps, and calculation of the credible sets.

The only missing piece to the IBSS-ss algorithm is the expression for the expected residual sum of squares (ERSS) under the variational approximation to the posterior, $q(\boldsymbol{b})$, which is needed to estimate $\sigma^2$. Again, the expression can be written in terms of the sufficient statistics:

$$\begin{aligned}
&\mathrm{ERSS}(\boldsymbol{X}, \boldsymbol{y}, \bar{\boldsymbol{b}}_1, \ldots, \bar{\boldsymbol{b}}_L, \bar{\boldsymbol{b}^2_1}, \ldots, \bar{\boldsymbol{b}^2_L}]) \\
&\quad := \mathbb{E}_q[\|\boldsymbol{y} - \boldsymbol{X}\boldsymbol{b}\|^2] \\
&\quad = \|\boldsymbol{y} - \boldsymbol{X}\bar{\boldsymbol{b}}\|^2 - \sum_{l=1}^{L} \bar{\boldsymbol{b}}_l \boldsymbol{X}^\intercal \boldsymbol{X} \bar{\boldsymbol{b}}_l + \sum_{l=1}^{L}\sum_{j=1}^{J} (\boldsymbol{x}_j^\intercal \boldsymbol{x}_j) \bar{b^2_{lj}} \\
&\quad = \boldsymbol{y}^\intercal \boldsymbol{y} - 2\bar{\boldsymbol{b}}^\intercal \boldsymbol{X}^\intercal \boldsymbol{y} + \bar{\boldsymbol{b}}^\intercal \boldsymbol{X}^\intercal \boldsymbol{X} \bar{\boldsymbol{b}} - \sum_{l=1}^{L} \bar{\boldsymbol{b}}_l^\intercal \boldsymbol{X}^\intercal \boldsymbol{X} \bar{\boldsymbol{b}}_l + \sum_{l=1}^{L}\sum_{j=1}^{J} (\boldsymbol{x}_j^\intercal \boldsymbol{x}_j) \bar{b^2_{lj}} \\
&\quad := \mathrm{ERSS\text{-}ss}(\boldsymbol{X}^\intercal \boldsymbol{X}, \boldsymbol{X}^\intercal \boldsymbol{y}, \boldsymbol{y}^\intercal \boldsymbol{y}, \bar{\boldsymbol{b}}_1, \ldots, \bar{\boldsymbol{b}}_L, \bar{\boldsymbol{b}^2_1}, \ldots, \bar{\boldsymbol{b}^2_L}). \qquad (41)
\end{aligned}$$

The second ERSS definition makes explicit that it is a function of the sufficient statistics.

#### Computing sufficient statistics, and approximations, from summary data

**Lemma 1** (Computing sufficient statistics)**.** The statistics $\boldsymbol{X}^\intercal\boldsymbol{X}, \boldsymbol{X}^\intercal\boldsymbol{y}$ can be computed from the the summary data $\hat{\boldsymbol{b}}, \hat{\boldsymbol{s}}, \boldsymbol{R}, \boldsymbol{y}^\intercal\boldsymbol{y}, N$ from the following formulae:

$$\hat{\sigma}_j^2 = \frac{\boldsymbol{y}^\intercal\boldsymbol{y}}{\hat{b}_j^2/\hat{s}_j^2 + N - 2} \qquad (42)$$

$$\boldsymbol{x}_j^\intercal \boldsymbol{y} = \hat{\sigma}_j^2 \hat{b}_j / \hat{s}_j^2 \qquad (43)$$

$$\boldsymbol{x}_j^\intercal \boldsymbol{x}_j = \hat{\sigma}_j^2 / \hat{s}_j^2 \qquad (44)$$

$$\boldsymbol{D}_{xx} = \mathrm{diag}(\boldsymbol{x}_1^\intercal \boldsymbol{x}_1, \ldots, \boldsymbol{x}_J^\intercal \boldsymbol{x}_J) \qquad (45)$$

$$\boldsymbol{X}^\intercal \boldsymbol{X} = \boldsymbol{D}_{xx}^{1/2} \boldsymbol{R} \boldsymbol{D}_{xx}^{1/2}. \qquad (46)$$

*Proof.* The correlation coefficient $r_j^2$ for a simple linear regression of $\boldsymbol{y}$ on SNP $j$ is:

$$r_j^2 := \frac{\hat{b}_j^2/\hat{s}_j^2}{\hat{b}_j^2/\hat{s}_j^2 + N - 2}. \qquad (47)$$

The estimated residual variance from the simple linear regression model is therefore

$$\begin{aligned}
\hat{\sigma}_j^2 &= \frac{\boldsymbol{y}^\intercal\boldsymbol{y}(1 - r_j^2)}{N - 2} \\
&= \frac{\boldsymbol{y}^\intercal\boldsymbol{y}}{\hat{b}_j^2/\hat{s}_j^2 + N - 2}. \qquad (48)
\end{aligned}$$

The remaining expressions follow straightforwardly from standard statistical formulas for the sample correlation and point estimation in linear regression. □

**Remark 2.** The formulae (42–46) are ordered in such a way that to provide a step-by-step procedure for reconstructing $\boldsymbol{X}^\intercal\boldsymbol{X}, \boldsymbol{X}^\intercal\boldsymbol{y}$ from the summary data $\hat{\boldsymbol{b}}, \hat{\boldsymbol{s}}, \boldsymbol{R}, \boldsymbol{y}^\intercal\boldsymbol{y}, N$. Thus, given $\hat{\boldsymbol{b}}, \hat{\boldsymbol{s}}, \boldsymbol{R}, \boldsymbol{y}^\intercal\boldsymbol{y}, N$, *XXX* involves first computing $\boldsymbol{X}^\intercal\boldsymbol{X}, \boldsymbol{X}^\intercal\boldsymbol{y}$ using (42–46), and then applying the IBSS-ss algorithm to $\boldsymbol{X}^\intercal\boldsymbol{X}, \boldsymbol{X}^\intercal\boldsymbol{y}, \boldsymbol{y}^\intercal\boldsymbol{y}, N$.

**Remark 3.** Often, the in-sample LD matrix $\boldsymbol{R}$ is not available, and is replaced with an estimate $\hat{\boldsymbol{R}}$. Given $\hat{\boldsymbol{b}}, \hat{\boldsymbol{s}}, \hat{\boldsymbol{R}}, \boldsymbol{y}^\intercal\boldsymbol{y}, N$, *SuSiE-RSS* involves first computing $\boldsymbol{V}_{xx}, \boldsymbol{X}^\intercal\boldsymbol{y}$ from (42–46), with $\hat{\boldsymbol{R}}$ replacing $\boldsymbol{R}$ in the formula for $\boldsymbol{X}^\intercal\boldsymbol{X}$ to yield $N\boldsymbol{V}_{xx}$, then applying the IBSS-ss algorithm to $N\boldsymbol{V}_{xx}, \boldsymbol{X}^\intercal\boldsymbol{y}$ with $\sigma^2 = \boldsymbol{y}^\intercal\boldsymbol{y}/N$. Note that when $\sigma^2$ is fixed, the IBSS-ss algorithm does not need $\boldsymbol{y}^\intercal\boldsymbol{y}$ or $N$.

##### Dealing with uncentered data

As stated above, $\boldsymbol{y}$ and the columns of $\boldsymbol{X}$ are assumed to be centered, and accordingly the sufficient statistics should be computed using a centered $\boldsymbol{X}, \boldsymbol{y}$. If the sufficient statistics have been computed from an $\boldsymbol{X}, \boldsymbol{y}$ that have not been centered, the sufficient statistics can be modified after the fact so that they correspond to a centered $\boldsymbol{X}$ and $\boldsymbol{y}$. Denoting the unmodified data as $\boldsymbol{X}, \boldsymbol{y}$ and the centered data as $\tilde{\boldsymbol{X}} := \boldsymbol{X} - \mathbf{1}_N \bar{\boldsymbol{x}}^\intercal, \tilde{\boldsymbol{y}} := \boldsymbol{y} - \bar{y}\mathbf{1}_N$, where $\bar{\boldsymbol{x}} = \boldsymbol{X}^\intercal \mathbf{1}_N / N$ is the vector of column means, $\bar{y} := \sum_{i=1}^N y_i$, and $\mathbf{1}_N$ is a column vector of ones of length $N$, the centering calculations for the sufficient statistics are

$$
\begin{aligned}
\tilde{\boldsymbol{X}}^\intercal \tilde{\boldsymbol{X}} &= \boldsymbol{X}^\intercal \boldsymbol{X} - N \bar{\boldsymbol{x}} \bar{\boldsymbol{x}}^\intercal \\
\tilde{\boldsymbol{X}}^\intercal \tilde{\boldsymbol{y}} &= \boldsymbol{X}^\intercal \boldsymbol{y} - N \bar{y} \bar{\boldsymbol{x}} \\
\tilde{\boldsymbol{y}}^\intercal \tilde{\boldsymbol{y}} &= \boldsymbol{y}^\intercal \boldsymbol{y} - N \bar{y}^2.
\end{aligned}
$$

Similarly, the sufficient statistics may also be modified *post hoc* to be as if they were computed using a column-standardized $\boldsymbol{X}$; that is, an $\boldsymbol{X}$ in which each column has unit variance. Denoting the standardized (and centered) matrix as $\hat{\boldsymbol{X}}$, and assuming $\boldsymbol{X}$ is centered, the calculations are

$$
\begin{aligned}
\hat{\boldsymbol{X}}^\intercal \hat{\boldsymbol{X}} &= (N-1) \times \boldsymbol{D}_{xx}^{-1/2} \boldsymbol{X}^\intercal \boldsymbol{X} \boldsymbol{D}_{xx}^{-1/2} \\
\hat{\boldsymbol{X}}^\intercal \boldsymbol{y} &= \sqrt{N-1} \times \boldsymbol{D}_{xx}^{-1/2} \boldsymbol{X}^\intercal \boldsymbol{y}.
\end{aligned}
$$

In these formulae we have assumed that unbiased sample variances ($N-1$ denominator) are used to standardize columns of $\boldsymbol{X}$.

#### Alternative model-based derivation

We derived *SuSiE-RSS* by writing down the full-data likelihood in terms of sufficient statistics, approximating the sufficient statistics, then substituting the approximations to obtain an approximate likelihood (12). In the main text, we mentioned that, in the standardized case, the resulting approximate likelihood can also be obtained from (22). Here we derive the more general version of this model when $\boldsymbol{X}, \boldsymbol{y}$ are not necessarily standardized.

Recall, the model for individual-level data is

$$\boldsymbol{y} \sim \mathcal{N}_J(\boldsymbol{X}\boldsymbol{b}, \sigma^2 \boldsymbol{I}_N). \tag{49}$$

From this, we have

$$\boldsymbol{X}^\intercal \boldsymbol{y}/\sqrt{N} \sim \mathcal{N}_J(\boldsymbol{X}^\intercal \boldsymbol{X}/N \times \sqrt{N}\boldsymbol{b}, \sigma^2 \boldsymbol{X}^\intercal \boldsymbol{X}/N). \tag{50}$$

If $\boldsymbol{X}^\intercal \boldsymbol{X}$ is invertible, the density obtained from (50) yields the same likelihood (up to a constant of proportionality) as the likelihood from the individual-level data (8). Substituting $\boldsymbol{X}^\intercal \boldsymbol{X}$ with $N\boldsymbol{V}_{xx}$ yields

$$\boldsymbol{X}^\intercal \boldsymbol{y}/\sqrt{N} \sim \mathcal{N}_J(\boldsymbol{V}_{xx} \times \sqrt{N}\boldsymbol{b}, \sigma^2 \boldsymbol{V}_{xx}). \tag{51}$$

If $\boldsymbol{V}_{xx}$ is invertible, the density obtained from (51) yields the same likelihood (up to a constant of proportionality) as (12).

In the special case where $\boldsymbol{X}$ and $\boldsymbol{y}$ are standardized, these expressions simplify. Making substitutions $\boldsymbol{X}^\intercal \boldsymbol{y}/\sqrt{N} = \tilde{\boldsymbol{z}}$ and $\boldsymbol{X}^\intercal \boldsymbol{X}/N = \boldsymbol{R}$ in (50) results in the simplified expression

$$\tilde{\boldsymbol{z}} \sim \mathcal{N}_J(\sqrt{N}\boldsymbol{R}\boldsymbol{b}, \sigma^2 \boldsymbol{R}). \tag{52}$$

Finally, substituting $\hat{\boldsymbol{R}}$ for $\boldsymbol{R}$ and fixing $\sigma^2 = 1$ gives (22).

#### Dealing with non-invertible LD matrix

If $\hat{\boldsymbol{R}}$ is not invertible, the models (20) and (22) do not have a density (with respect to Lebesgue measure). This complicates defining a likelihood from these models. Various approaches have been proposed for dealing with this issue. Some of these approaches are equivalent to using the likelihood (19) that we use here, which exists whether or not $\hat{\boldsymbol{R}}$ is invertible, while other approaches are not equivalent. Here we argue that using the likelihood (19) has the advantage of being simple, and satisfies the desirable property that inference under the SER model is independent of LD. We call this property "Irrelevance of null SNPs."

Note that the issues discussed here remain the same whether one uses models and likelihoods for the observed $z$-scores, $\hat{\boldsymbol{z}}$, or the PVE-adjusted $z$-scores, $\tilde{\boldsymbol{z}}$. Therefore, to simplify presentation we describe the results for the $z$-scores, noting that one can substitute for the PVE-adjusted $z$-scores wherever $\hat{\boldsymbol{z}}$ appears in the mathematical expressions below. To further simplify, we use $\boldsymbol{z} := \sqrt{N}\boldsymbol{b}$ to denote the scaled parameters, and compare the model

$$\hat{\boldsymbol{z}} \sim \mathcal{N}_J(\hat{\boldsymbol{R}}\boldsymbol{z}, \hat{\boldsymbol{R}}) \tag{53}$$

and the likelihood

$$\ell_{\mathit{RSS\text{-}Z}}(\boldsymbol{z}) := \exp(-\tfrac{1}{2}\boldsymbol{z}^\intercal \hat{\boldsymbol{R}}\boldsymbol{z} + \boldsymbol{z}^\intercal \hat{\boldsymbol{z}}), \tag{54}$$

which is the same (up to a constant of proportionality) as (19) but with $\hat{\boldsymbol{z}}$ replacing $\tilde{\boldsymbol{z}}$ and $\boldsymbol{z}$ replacing $\sqrt{N}\boldsymbol{b}$. If $\hat{\boldsymbol{R}}$ is invertible, the density of the model (53) yields likelihood (54). We call (53) the "*RSS-Z* model" and (54) the "*RSS-Z* likelihood". The key point of this section is to argue that under model (53) the likelihood (54) is the "correct" likelihood even if $\hat{\boldsymbol{R}}$ is not invertible.

We assume that although $\hat{\boldsymbol{R}}$ may not be invertible it is nonetheless a valid covariance matrix. That is, it is symmetric and positive semidefinite (PSD), which means that all its eigenvalues are non-negative. This is guaranteed so long as $\hat{\boldsymbol{R}}$ is a sample correlation matrix. However, it may be violated if $\hat{\boldsymbol{R}}$ is obtained by modifying a sample correlation matrix, for example by setting small correlations to zero. Any $J \times J$ symmetric PSD matrix $\hat{\boldsymbol{R}}$ with rank $r \leq J$ has an eigenvalue decomposition of the form

$$\hat{\boldsymbol{R}} = \boldsymbol{Q}\boldsymbol{\Lambda}\boldsymbol{Q}^\intercal, \tag{55}$$

where $\boldsymbol{\Lambda}$ is an $r \times r$ diagonal matrix with the $r$ positive eigenvalues $\lambda_1 \geq \lambda_2 \cdots \geq \lambda_r > 0$ along its diagonal, and $\boldsymbol{Q}$ is a $J \times r$ matrix whose columns are the $r$ eigenvectors corresponding to the $r$ non-zero eigenvalues, and $\boldsymbol{Q}^\intercal\boldsymbol{Q} = \boldsymbol{I}_r$.

One can modify $\hat{\boldsymbol{R}}$ to make it invertible by simply adding a small diagonal element. Indeed, for any $\lambda \in (0, 1)$, the matrix

$$\hat{\boldsymbol{R}}_\lambda := (1 - \lambda)\hat{\boldsymbol{R}} + \lambda\boldsymbol{I} \tag{56}$$

will be invertible.

When $\hat{\boldsymbol{R}}$ is not invertible, the *RSS-Z* model (53) becomes degenerate, which means that some values of $\hat{\boldsymbol{z}}$ become impossible. In particular, with probability one, $\hat{\boldsymbol{z}} \in \text{range}(\boldsymbol{Q})$; that is, $\hat{\boldsymbol{z}} = \boldsymbol{Q}\boldsymbol{\alpha}$ for some $\boldsymbol{\alpha}$.

**Definition 1** (Consistency of $\hat{\boldsymbol{z}}$ with $\hat{\boldsymbol{R}}$)**.** We say that $\hat{\boldsymbol{z}}$ is *consistent* with $\hat{\boldsymbol{R}}$ if $\hat{\boldsymbol{z}} \in \text{range}(\boldsymbol{Q})$. Otherwise, if $\hat{\boldsymbol{z}} \notin \text{range}(\boldsymbol{Q})$ we say $\hat{\boldsymbol{z}}$ is *inconsistent* with $\hat{\boldsymbol{R}}$.

Note that, if $\hat{\boldsymbol{R}}$ is invertible, then $\text{range}(\boldsymbol{Q}) = \mathbb{R}^J$, and so $\hat{\boldsymbol{z}}$ will always be consistent with $\hat{\boldsymbol{R}}$. Further, if $\hat{\boldsymbol{z}}$ was generated from the model (53), then it will be consistent with $\hat{\boldsymbol{R}}$ (with probability 1). However, in practice the model (53) is only an approximation, and so in practice $\hat{\boldsymbol{z}}$ may be inconsistent with $\hat{\boldsymbol{R}}$. (Even when $\hat{\boldsymbol{R}} = \boldsymbol{R}$, $\hat{\boldsymbol{z}}$ could be inconsistent with $\hat{\boldsymbol{R}}$; this is not true for $\tilde{\boldsymbol{z}}$.)

Now we consider four approaches to dealing with a non-invertible $\hat{\boldsymbol{R}}$.

1a. Substitute the non-invertible $\hat{\boldsymbol{R}}$ with the invertible matrix $\hat{\boldsymbol{R}}_\lambda$, for some small $\lambda$, in (53). This is the approach used in [4,5]. The model becomes $\hat{\boldsymbol{z}} \sim \mathcal{N}_J(\hat{\boldsymbol{R}}_\lambda \boldsymbol{z}, \hat{\boldsymbol{R}}_\lambda)$, which has a density because $\hat{\boldsymbol{R}}_\lambda$ is invertible, yielding likelihood

$$\ell_{1a}(\boldsymbol{z}; \hat{\boldsymbol{z}}, \hat{\boldsymbol{R}}, \lambda) := \exp(-\tfrac{1}{2}\boldsymbol{z}^\intercal \hat{\boldsymbol{R}}_\lambda \boldsymbol{z} + \boldsymbol{z}^\intercal \hat{\boldsymbol{z}}). \tag{57}$$

1b. Use the *RSS-Z* likelihood (54) even though $\hat{\boldsymbol{R}}$ is non-invertible, so the likelihood is

$$\ell_{1b}(\boldsymbol{z}; \hat{\boldsymbol{z}}, \hat{\boldsymbol{R}}) := \exp(-\tfrac{1}{2}\boldsymbol{z}^\intercal \hat{\boldsymbol{R}} \boldsymbol{z} + \boldsymbol{z}^\intercal \hat{\boldsymbol{z}}). \tag{58}$$

Note that the likelihood exists, and is easily computed, even when $\hat{\boldsymbol{R}}$ is not invertible. Earlier versions of FINEMAP [6] effectively used this approach (disallowing configurations $\gamma \subseteq \{1, \ldots, J\}$ where $\hat{\boldsymbol{R}}_\gamma$ is not invertible).

2a. Set the covariance in the *RSS-Z* model (53) to $\hat{\boldsymbol{R}}_\lambda$ for some small $\lambda$, so that $\hat{\boldsymbol{z}} \sim \mathcal{N}_J(\hat{\boldsymbol{R}}\boldsymbol{z}, \hat{\boldsymbol{R}}_\lambda)$. Note that this approach differs from 1a because it uses $\hat{\boldsymbol{R}}$ instead of $\hat{\boldsymbol{R}}_\lambda$ for the mean. This yields the following likelihood:

$$\ell_{2a}(\boldsymbol{z}; \hat{\boldsymbol{z}}, \hat{\boldsymbol{R}}, \lambda) := \exp(-\tfrac{1}{2}\boldsymbol{z}^\intercal \hat{\boldsymbol{R}}\hat{\boldsymbol{R}}_\lambda^{-1}\hat{\boldsymbol{R}} \boldsymbol{z} + \boldsymbol{z}^\intercal \hat{\boldsymbol{R}}\hat{\boldsymbol{R}}_\lambda^{-1} \hat{\boldsymbol{z}}). \tag{59}$$

2b. Project $\hat{\boldsymbol{z}}$ into a lower-dimensional subspace, $\hat{\boldsymbol{z}}' := \boldsymbol{\Lambda}^{-1/2}\boldsymbol{Q}^\intercal \hat{\boldsymbol{z}}$, which ensures that $\hat{\boldsymbol{z}}' \in \mathbb{R}^r$ has a probability density, $\hat{\boldsymbol{z}}' \sim \mathcal{N}_r(\boldsymbol{\Lambda}^{1/2}\boldsymbol{Q}^\intercal \boldsymbol{z}, \boldsymbol{I}_r)$. Thus we have

$$\begin{aligned} p(\hat{\boldsymbol{z}}' \mid \boldsymbol{z}, \hat{\boldsymbol{R}}) &\propto \exp\{-\tfrac{1}{2}(\hat{\boldsymbol{z}}' - \boldsymbol{\Lambda}^{1/2}\boldsymbol{Q}^\intercal \boldsymbol{z})^\intercal (\hat{\boldsymbol{z}}' - \boldsymbol{\Lambda}^{1/2}\boldsymbol{Q}^\intercal \boldsymbol{z})\} \\ &\propto \exp(-\tfrac{1}{2}\boldsymbol{z}^\intercal \hat{\boldsymbol{R}} \boldsymbol{z} + \boldsymbol{z}^\intercal \boldsymbol{Q}\boldsymbol{\Lambda}^{1/2} \hat{\boldsymbol{z}}') \\ &= \exp(-\tfrac{1}{2}\boldsymbol{z}^\intercal \hat{\boldsymbol{R}} \boldsymbol{z} + \boldsymbol{z}^\intercal \boldsymbol{Q}\boldsymbol{Q}^\intercal \hat{\boldsymbol{z}}), \end{aligned} \tag{60}$$

and therefore the likelihood is

$$\ell_{2b}(\boldsymbol{z}; \hat{\boldsymbol{z}}, \hat{\boldsymbol{R}}) = \exp(-\tfrac{1}{2}\boldsymbol{z}^\intercal \hat{\boldsymbol{R}} \boldsymbol{z} + \boldsymbol{z}^\intercal \boldsymbol{Q}\boldsymbol{Q}^\intercal \hat{\boldsymbol{z}}). \tag{61}$$

The same likelihood is obtained by replacing $\hat{\boldsymbol{R}}^{-1}$ in the density for (53) with the pseudoinverse of $\hat{\boldsymbol{R}}$, which is $\boldsymbol{Q}\boldsymbol{\Lambda}^{-1}\boldsymbol{Q}^\intercal$. This is the approach used in msCAVIAR [7].

We summarize the connections between these four approaches in the following proposition.

**Proposition 2.** (a) As $\lambda \to 0$, Approach 1a becomes equivalent to Approach 1b, and Approach 2a becomes equivalent to Approach 2b; that is,

$$\lim_{\lambda \to 0} \ell_{1a}(\boldsymbol{z}; \hat{\boldsymbol{z}}, \hat{\boldsymbol{R}}, \lambda) = \ell_{1b}(\boldsymbol{z}; \hat{\boldsymbol{z}}, \hat{\boldsymbol{R}}); \tag{62}$$

$$\lim_{\lambda \to 0} \ell_{2a}(\boldsymbol{z}; \hat{\boldsymbol{z}}, \hat{\boldsymbol{R}}, \lambda) = \ell_{2b}(\boldsymbol{z}; \hat{\boldsymbol{z}}, \hat{\boldsymbol{R}}). \tag{63}$$

(b) Approaches 1b and 2b are equivalent—*i.e.*, $\ell_{1b}(\boldsymbol{z}; \hat{\boldsymbol{z}}, \hat{\boldsymbol{R}}) = \ell_{2b}(\boldsymbol{z}; \hat{\boldsymbol{z}}, \hat{\boldsymbol{R}})$—if and only if $\hat{\boldsymbol{z}}$ is consistent with $\hat{\boldsymbol{R}}$.

(c) If $\hat{\boldsymbol{z}}$ is inconsistent with $\hat{\boldsymbol{R}}$, Approach 2b behaves discontinuously,

$$\lim_{\lambda \to 0} \ell_{2b}(\boldsymbol{z}; \hat{\boldsymbol{z}}, \hat{\boldsymbol{R}}_\lambda) \neq \ell_{2b}(\boldsymbol{z}; \hat{\boldsymbol{z}}, \hat{\boldsymbol{R}}), \tag{64}$$

but in the limit it is equivalent to Approach 1b,

$$\lim_{\lambda \to 0} \ell_{2b}(\boldsymbol{z}; \hat{\boldsymbol{z}}, \hat{\boldsymbol{R}}_\lambda) = \ell_{1b}(\boldsymbol{z}; \hat{\boldsymbol{z}}, \hat{\boldsymbol{R}}). \tag{65}$$

*Proof.* (a) As $\lambda \to 0$, $\hat{\boldsymbol{R}}_\lambda \to \hat{\boldsymbol{R}}$, so (62) is trivially satisfied. To prove (63), we define $\boldsymbol{B} := \boldsymbol{Q}\boldsymbol{\Lambda}^{1/2}$ with pseudoinverse $\boldsymbol{B}^\dagger = \boldsymbol{\Lambda}^{-1/2}\boldsymbol{Q}^\intercal$. With these definitions, we can write $\ell_{2a}(\boldsymbol{z}; \hat{\boldsymbol{z}}, \hat{\boldsymbol{R}}, \lambda)$ as

$$\begin{aligned}\ell_{2a}(\boldsymbol{z}; \hat{\boldsymbol{z}}, \hat{\boldsymbol{R}}, \lambda) = \exp(-\tfrac{1}{2}\boldsymbol{z}^\intercal \boldsymbol{B}\boldsymbol{B}^\intercal((1-\lambda)\boldsymbol{B}\boldsymbol{B}^\intercal + \lambda \boldsymbol{I})^{-1}\boldsymbol{B}\boldsymbol{B}^\intercal \boldsymbol{z} \\ + \boldsymbol{z}^\intercal \boldsymbol{B}\boldsymbol{B}^\intercal((1-\lambda)\boldsymbol{B}\boldsymbol{B}^\intercal + \lambda \boldsymbol{I})^{-1}\hat{\boldsymbol{z}}).\end{aligned} \tag{66}$$

In the limit as $\lambda \to 0$, $\boldsymbol{B}^\intercal((1-\lambda)\boldsymbol{B}\boldsymbol{B}^\intercal + \lambda\boldsymbol{I})^{-1} \to \boldsymbol{B}^\dagger$ (Theorem 3.4 in [8]). Therefore,

$$\begin{aligned}\lim_{\lambda\to 0} \ell_{2a}(\boldsymbol{z}; \hat{\boldsymbol{z}}, \hat{\boldsymbol{R}}, \lambda) &= \exp(-\tfrac{1}{2}\boldsymbol{z}^\intercal \boldsymbol{B}\boldsymbol{B}^\dagger\boldsymbol{B}\boldsymbol{B}^\intercal \boldsymbol{z} + \boldsymbol{z}^\intercal \boldsymbol{B}\boldsymbol{B}^\dagger \hat{\boldsymbol{z}}) \\ &= \exp(-\tfrac{1}{2}\boldsymbol{z}^\intercal \hat{\boldsymbol{R}} \boldsymbol{z} + \boldsymbol{z}^\intercal \boldsymbol{Q}\boldsymbol{Q}^\intercal \hat{\boldsymbol{z}}) \\ &= \ell_{2b}(\boldsymbol{z}; \hat{\boldsymbol{z}}, \hat{\boldsymbol{R}}).\end{aligned} \tag{67}$$

(b) $\ell_{1b}(\boldsymbol{z}; \hat{\boldsymbol{z}}, \hat{\boldsymbol{R}}) = \ell_{2b}(\boldsymbol{z}; \hat{\boldsymbol{z}}, \hat{\boldsymbol{R}})$ for all $\boldsymbol{z}$ if and only if $\boldsymbol{Q}\boldsymbol{Q}^\intercal \hat{\boldsymbol{z}} = \hat{\boldsymbol{z}}$. Since $\boldsymbol{P} = \boldsymbol{Q}\boldsymbol{Q}^\intercal$ is an orthogonal projector onto range($\boldsymbol{Q}$), $\hat{\boldsymbol{z}} \in \text{range}(Q) \Leftrightarrow \boldsymbol{Q}\boldsymbol{Q}^\intercal \hat{\boldsymbol{z}} = \hat{\boldsymbol{z}}$ (see [9], Ch. 6).

(c) First we prove (65). Since $\hat{\boldsymbol{R}}_\lambda$ is full rank, the $J \times J$ matrix of eigenvectors $\boldsymbol{Q}_\lambda$ satisfies $\boldsymbol{Q}_\lambda \boldsymbol{Q}_\lambda^\intercal = \boldsymbol{I}_J$. Therefore,

$$\ell_{2b}(\boldsymbol{z}; \hat{\boldsymbol{z}}, \hat{\boldsymbol{R}}_\lambda) = \exp(-\tfrac{1}{2}\boldsymbol{z}^\intercal \hat{\boldsymbol{R}}_\lambda \boldsymbol{z} + \boldsymbol{z}^\intercal \hat{\boldsymbol{z}}) = \ell_{1a}(\boldsymbol{z}; \hat{\boldsymbol{z}}, \hat{\boldsymbol{R}}_\lambda), \tag{68}$$

so the result follows from (62). The result (64) then follows from part (b). □

**Remark 4.** Proposition 2 (a) and (b) together imply that if $\hat{\boldsymbol{z}}$ is consistent with $\hat{\boldsymbol{R}}$ then, for sufficiently small $\lambda$, all four approaches should give the same results (ignoring numerical errors that occur in floating-point computations). However, because the *RSS-Z* model is only an approximation, $\hat{\boldsymbol{z}}$ will often be inconsistent with $\hat{\boldsymbol{R}}$. It is this fact that causes the methods to give different results.

##### Irrelevance of null SNPs

Given that the different approaches to dealing with non-invertible $\hat{\boldsymbol{R}}$ may give different results, it is natural to ask which approach is preferable. Here we argue that Approaches 1a and 1b are preferable to Approaches 2a and 2b because Approaches 1a and 1b always satisfy a simple property that we call "irrelevance of null SNPs." (Approaches 2a and 2b may sometimes satisfy this property, but they are not guaranteed to do so.) This property is also satisifed by the full data likelihood and implies, among other things, that inference under the SER model is independent of the LD matrix.

We motivate this property by observing that the individual-data multiple regression likelihood (1) has the following simple property: if a SNP $j$ has no effect (a "null SNP"), then its genotypes $\boldsymbol{x}_j$ do not appear in the likelihood. As a result, genotypes at null SNPs do not impact inference for other SNPs. We call this property "irrelevance of null SNPs."

In the summary-data setting, we do not directly observe the genotypes, so we need to formulate an analogous property. Therefore, to translate these ideas to the summary-data likelihoods with $\boldsymbol{z}, \hat{\boldsymbol{R}}$, we ask instead whether the likelihood has the following property: if $z_j = 0$, then $\hat{R}_{j1}, \ldots, \hat{R}_{jJ}$ do not appear in the likelihood. If, for example, $\hat{R}_{jj'}$ is the correlation between SNPs $j$ and $j'$ obtained from a suitable reference panel, then this property implies that the genotypes for SNP $j$ have no impact on the summary-data likelihood when $z_j = 0$. (The same can be said for regularized LD matrices of the form (23).)

To formalize these ideas, we introduce some notation. We use $\gamma$ to denote a subset of SNPs, $\gamma \subseteq \{1, \ldots, J\}$, and we let $\boldsymbol{z}_\gamma$ denote the elements of the vector $\boldsymbol{z}$ corresponding to the SNPs in $\gamma$. The remaining elements are denoted by $\boldsymbol{z}_{-\gamma}$. Similarly, we use $\hat{\boldsymbol{R}}_\gamma$ to denote the $|\gamma| \times |\gamma|$ matrix formed by the rows and columns of $\hat{\boldsymbol{R}}$ corresponding to elements of $\gamma$, and $\hat{\boldsymbol{R}}_{-\gamma}$ denotes the matrix formed by all rows and columns of $\hat{\boldsymbol{R}}$ that do not correspond to elements of $\gamma$. We then define irrelevance of null SNPs as follows.

**Definition 2** (Irrelevance of null SNPs)**.** Let $\ell(\boldsymbol{z})$ be any likelihood for $\boldsymbol{z}$ (which implicitly depends on $\hat{\boldsymbol{z}}, \hat{\boldsymbol{R}}$, and the model parameters). For any subset $\gamma \subseteq \{1, \ldots, J\}$, let $\ell^\gamma(\boldsymbol{z}_\gamma)$ denote the likelihood for $\boldsymbol{z}_\gamma$ when the remaining elements $\boldsymbol{z}_{-\gamma}$ are set to zero; that is,

$$\ell^\gamma(\boldsymbol{z}_\gamma) := \ell(\boldsymbol{z}_\gamma, \boldsymbol{z}_{-\gamma} = \boldsymbol{0}). \tag{69}$$

We say the likelihood $\ell(\boldsymbol{z})$ satisfies the irrelevance of null SNPs property if, for all $\gamma$, $\ell^\gamma(\boldsymbol{z}_\gamma)$ depends on $\hat{\boldsymbol{R}}$ only through $\hat{\boldsymbol{R}}_\gamma$.

**Remark 5.** We have framed the definition in terms of likelihoods for the scaled parameters $\boldsymbol{z}$, but a similar definition could be obtained for the effects $\boldsymbol{b}$ by replacing $\boldsymbol{z}$ with $\boldsymbol{b}$.

The multiple regression likelihood based on individual-level data (8) satisfies the irrelevance of null SNPs property. Likelihoods $\ell_{1a}$ (57) and $\ell_{1b}$ (58) also satisfy this property, as summarized by the following proposition.

**Proposition 3.** *$\ell_{RSS\text{-}Z}$ (54) satisifies irrelevance of null SNPs (Definition 2).*

*Proof.* Setting $\boldsymbol{z}_{-\gamma} = \boldsymbol{0}$ in (54) yields

$$\ell_{RSS\text{-}Z}(\boldsymbol{z}_\gamma, \boldsymbol{z}_{-\gamma} = \boldsymbol{0}) = \exp(-\tfrac{1}{2}\boldsymbol{z}_\gamma^\intercal \hat{\boldsymbol{R}}_\gamma \boldsymbol{z}_\gamma + \boldsymbol{z}_\gamma^\intercal \hat{\boldsymbol{z}}_\gamma). \tag{70}$$

□

The irrelevance of null SNPs has the following simple implication: to assess support for the hypothesis $H_\gamma : \boldsymbol{z}_\gamma = 0$, one only needs the genotypes corresponding to the non-null SNPs. Indeed, if $p_\gamma(\boldsymbol{z}_\gamma)$ denotes any prior on $\boldsymbol{z}_\gamma$ under $H_\gamma$ then, in the absence of nuisance parameters, the Bayes Factor for $H_\gamma$ vs. $H_0$ is

$$\mathrm{BF}_\gamma := \frac{\int \ell^\gamma(\boldsymbol{z}_\gamma)\, p_\gamma(\boldsymbol{z}_\gamma)\, d\boldsymbol{z}_\gamma}{\ell^\gamma(\boldsymbol{z}_\gamma = \boldsymbol{0})}. \tag{71}$$

This result implies that $\mathrm{BF}_\gamma$ depends only on $\hat{\boldsymbol{z}}_\gamma, \hat{\boldsymbol{R}}_\gamma$, whatever priors are used for each $\gamma$ (assuming that the prior $p_\gamma$ does not depend on the null genotypes). This result is easily extended to integrate out additional nuisance parameters (e.g., $\sigma$ in the multiple regression model) in both the numerator and denominator.

A similar result is shown for specific priors in [6], and is exploited in FINEMAP. Our analysis here emphasizes that this is, fundamentally, due to properties of the likelihood, and is not confined to specific priors.

**Remark 6.** Applying this result to the special case that $\gamma$ contains a single SNP $j$, *i.e.*, $\gamma = \{j\}$, $\mathrm{BF}_\gamma$ depends on the genotypes only through SNP $j$; in particular, $\mathrm{BF}_\gamma$ *does not depend on the LD between SNPs.* Thus, irrelevance of null SNPs implies that fitting a SER does not depend on LD. (A similar observation has been made for propsective models of case-control traits, based on conditional-independence arguments [10]). Since the likelihood (13) satisfies irrelevance of null SNPs, inference under the SER model with this likelihood has the desirable property that it does not depend on LD.

#### Estimation of $\lambda$ in regularized LD matrix

To solve (24), we used the Brent-Dekker algorithm [11], which is implemented in R by the `optimize` function. This algorithm performs a 1-d search over $\lambda \in [0, 1]$. The main computational expense of this optimization step is the eigenvalue decomposition of $\hat{\boldsymbol{R}}_0$. Computing the eigenvalue decomposition has computational complexity $O(J^3)$, and therefore can impose a substantial computational burden on the overall fine-mapping analysis when $J$ is large. In practice, we found that the regularization typically provided only a small improvement to the *SuSiE* fine-mapping results, so in the software we set $\lambda = 0$ by default to avoid this potentially large computational expense.

#### Likelihood ratio for detecting allele flips

Based on the conditional distribution (26) and the standardized differences (27), we developed a likelihood ratio to detect allele flips. When $\hat{\boldsymbol{R}} = \boldsymbol{R}$, $t_j$ should be approximately standard normal; $t_j \sim \mathcal{N}(0, 1)$. However, when $\hat{\boldsymbol{R}}$ is estimated from a reference panel, even without errors such as allele flips, the empirical distribution of standardized differences will be longer tailed than the standard normal. This suggests that a more flexible distribution should be used to model the standardized differences. We used a mixture of normals to model the empirical conditional distribution,

$$\tilde{z}_j \mid \tilde{\boldsymbol{z}}_{-j}, \hat{\boldsymbol{R}} \sim \sum_{k=1}^{K} w_k \mathcal{N}(-\boldsymbol{\Omega}_{j,-j}\tilde{\boldsymbol{z}}_{-j}/\Omega_{jj}, \sigma_k^2/\Omega_{jj}), \tag{72}$$

where $\sigma_1, \ldots, \sigma_K$ are prespecified standard deviations such that $\sigma_1 < \cdots < \sigma_K$, and $\boldsymbol{w} = (w_1, \ldots, w_K)$ are mixture proportions (that is, they are all non-negative and sum to 1). In our analyses, we chose the $\sigma_k$'s such that $\sigma_1 = 0.8$, $\sigma_K = 2 \times \max\{|t_1|, \ldots, |t_J|\}$ and $\sigma_{k+1} = 1.05 \times \sigma_k$. We estimated $\boldsymbol{w}$ by maximum likelihood, using summary data for all SNPs. Computing the maximum-likelihood estimate of $\boldsymbol{w}$ is a convex optimization problem and can be solved efficiently using `mixsqp` [12]. (The effort involved in solving this convex optimization problem is negligible compared to inverting or factorizing $\hat{\boldsymbol{R}}$ when $J$ is large.) We then used the maximum-likelihood estimates of the mixture weights, $\hat{\boldsymbol{w}} = (\hat{w}_1, \ldots, \hat{w}_K)$, to compute a likelihood ratio for each SNP $j$,

$$\mathrm{LR}_j := \frac{\sum_{k=1}^{K} \hat{\omega}_k \mathcal{N}(\tilde{z}_j; \boldsymbol{\Omega}_{j,-j}\tilde{\boldsymbol{z}}_{-j}/\Omega_{jj}, \sigma_k^2/\Omega_{jj})}{\sum_{k=1}^{K} \hat{\omega}_k \mathcal{N}(\tilde{z}_j; -\boldsymbol{\Omega}_{j,-j}\tilde{\boldsymbol{z}}_{-j}/\Omega_{jj}, \sigma_k^2/\Omega_{jj})}. \tag{73}$$

This likelihood ratio can only identify errors in SNPs $j$ with $z$-scores that are large in magnitude. Therefore, after estimating the mixture weights, we focus on SNPs $j$ with $|\tilde{z}_j| > 2$; SNPs $j$ with $|\tilde{z}_j| > 2$ and the largest likelihood ratios $\mathrm{LR}_j$ are the top candidates for being allele flips.

To verify the use of this likelihood ratio to identify allele flips, we simulated fine-mapping data sets in which exactly one SNP was a casual SNP, and exactly one of the SNPs had a flipped allele—that is, the allele encoding used to compute the $z$-scores was different from of the allele encoding used to compute the LD matrix. We considered two scenarios in these simulations: in the first scenario, the allele-flip SNP was also the causal SNP; in the second scenario, the allele-flip SNP was not the causal SNP. An allele flip in both of these scenarios can affect accuracy of the fine-mapping so it is important to identify and eliminate the allele-flip SNP in both cases.

We simulated fine-mapping data sets using the UK Biobank genotypes, as previously described in the Results (see also "Details of simulations" below), with the following changes: we simulated one causal SNP with an effect size chosen so that it explained 2% of variance in $\boldsymbol{y}$; and we used $N = 10{,}000$ samples to compute the $z$-scores and LD matrix. The $z$-scores and LD matrix were computed using the same samples ("in-sample LD"), except that the LD matrix was computed only after modifying the genotypes of the allele-flip SNP. Once the summary data $\hat{\boldsymbol{z}}, \hat{\boldsymbol{R}}$ were computed from the individual-level data, we computed $\mathrm{LR}_j$ for each SNP $j$. (Note that the likelihood ratios were computed using the unadjusted $z$-scores, $\hat{z}_j$, instead of the PVE-adjusted $z$-scores, $\tilde{z}_j$. The latter is recommended, but here this would have made little difference because the sample size was large and the SNP effects were small.) In total, we simulated 200 data sets for the first scenario and another 200 data sets for the second scenario.

The results of these simulations are summarized in S1 Fig. The allele-flip SNPs almost always had a likelihood ratio greater than 1 (*i.e.,* $\log \mathrm{LR}_j > 0$), and other SNPs usually had likelihood ratios less than 1 (note the logarithmic scale in some of these plots). While a small proportion of SNPs without an allele flip had likelihood ratios greater than 1 (plots in middle row), most of these SNPs had $z$-scores close to zero, so restricting to SNPs with larger $z$-scores (plots in bottom row) eliminates most of these larger likelihood ratios in non-allele-flip SNPs. In summary, the likelihood-ratio statistic (73) provides an accurate diagnostic for identification of allele flips so long as we restrict attention to SNPs with larger $z$-scores.

#### *SuSiE* refinement procedure

The refinement procedure is outlined in Algorithm 2. This procedure will work with any individual-level data or summary data accepted by *SuSiE*. In the algorithm, when we say the *SuSiE* model fit $s$ is initialized to fit $t$, specifically we mean that the posterior means $\bar{\boldsymbol{b}}_l$ for each single effect $l = 1, \ldots, L$ are initialized from $t$. (Refer to Algorithm 1 in [1], and Algorithm 1 in this paper.) The default initialization is $\bar{\boldsymbol{b}}_l = 0$. In all cases the default initialization is used for $\sigma^2$ and $\boldsymbol{\sigma_0^2} = (\sigma_{01}^2, \ldots, \sigma_{0L}^2)$.

**Algorithm 2** *SuSiE* refinement procedure

**Require:** A *SuSiE* model fit, $s$, with $K \geq 1$ credible sets, $CS_1, \ldots, CS_K$.

Compute ELBO at $s$, $F \leftarrow \mathrm{ELBO}(s)$
**repeat**
    **for** $k = 1$ to $K$ **do**
        $\tilde{\boldsymbol{\pi}} \leftarrow \boldsymbol{\pi}$
        Set prior weights to 0 for all SNPs in $CS_k$; $\tilde{\pi}_j \leftarrow 0$ for all $j \in CS_k$
        Fit *SuSiE* model, $t_k$, using prior weights $\tilde{\boldsymbol{\pi}}$ and default initialization
        Fit *SuSiE* model, $s_k$, using prior weights $\boldsymbol{\pi}$, initialized at $t_k$
        Compute ELBO at $s_k$, $F_k \leftarrow \mathrm{ELBO}(s_k)$
    **end for**
    $k^\star \leftarrow \mathrm{argmax}_k \, \mathrm{ELBO}(s_k)$
    **if** $F_{k^\star} > F$ **then**
        $s \leftarrow s_{k^\star}$
    **end if**
**until** $F_{k^\star} \leq F$

#### Details of calculations for toy example

In the toy example (see "Fine-mapping with inconsistent summary data and a non-invertible LD matrix: an illustration" in the Results), we assumed $\hat{\boldsymbol{R}}$ is the $2 \times 2$ rank-1 matrix of all ones. Then the eigenvalue decomposition of $\hat{\boldsymbol{R}}$ is $\hat{\boldsymbol{R}} = \boldsymbol{Q}\boldsymbol{\Lambda}\boldsymbol{Q}^\intercal$ with $\boldsymbol{\Lambda} = 1$, $\boldsymbol{Q} = (\sqrt{1/2}, \sqrt{1/2})^\intercal$, and $\boldsymbol{Q}\boldsymbol{Q}^T$ is the $2 \times 2$ matrix with all entries set to 1/2. In the likelihood $\ell_{2b}(\boldsymbol{z}; \hat{\boldsymbol{z}}, \hat{\boldsymbol{R}})$ (see eq. 61), this has the effect that $\boldsymbol{Q}\boldsymbol{Q}^\intercal\hat{\boldsymbol{z}}$ is the average of the observed $z$-scores; that is, $\boldsymbol{Q}\boldsymbol{Q}^\intercal\hat{\boldsymbol{z}} = ((\hat{z}_1 + \hat{z}_2)/2, (\hat{z}_1 + \hat{z}_2)/2)^\intercal$.

The *SuSiE-RSS* results for this toy example with $\hat{\boldsymbol{z}} = (6, 7)$ were generated by running `susie_rss` with the default settings (`susieR` version 0.11.48).

#### Details of simulations

We evaluated the fine-mapping methods on summary data sets generated using real genotypes and simulated phenotypes. For genotype data, we used version 3 of the imputed genotypes from the UK Biobank resource [13]. UK Biobank is a large-scale biomedical database and research resource containing genetic, lifestyle and health information from half a million UK participants. The UK Biobank genotype data are well suited for fine-mapping because of their large sample size (approximately 500,000 genotyped individuals) and the high density of available genetic variants after genotype imputation [14].

Following [15], we took steps to filter out genotype samples, resulting in a candidate set of 274,549 samples. In detail, to limit confounding due to population structure, we considered only genotype samples marked as "White British" (based on a principal components analysis of the genotypes [14]). We then removed any samples from the "White British" subset that met one or more of the following criteria: mismatch between self-reported and genetic sex; outlier based on heterozygosity and/or rate of missing genotypes; has at least one close relative in the same data set (based on the UK Biobank's kinship and "relatedness" calculations); or does not have a measurement of standing height.

From this collection of 274,549 genotype samples, we chose uniformly at a random a subset of 50,000 samples, then we used this random subset of genotypes to generate a collection of genotypes for fine-mapping. We repeated this 200 times to generate 200 genotype data sets for fine-mapping. Each of these 200 data sets contained SNPs within selected regions on autosomal chromosomes. These regions contained roughly 1,000 SNPs, and were chosen so that no two regions contained the same SNPs. A SNP was included in a data set only if it satisfied all of the following criteria: SNP with at most two alleles; minor allele frequency of 1% or greater; and information score, which quantifies imputation quality, of 0.9 or greater. On average, 998 SNPs were included in a region. The smallest region contained 998 SNPs, and the largest contained 1,001 SNPs. The average size of a region in base pairs was 390 kb.

For each of the 200 randomly chosen regions, we generated three data sets by following a procedure similar to [1], resulting in a total of $200 \times 3 = 600$ fine-mapping data sets. Our procedure is briefly described here. We simulated phenotypes $\boldsymbol{y}$ under the multiple regression model (1) in which $\mathbf{X}$ was the centered and standardized matrix of 50,000 genotypes. We simulated three sets of phenotype data from the same $\mathbf{X}$ by setting the number of causal SNPs to be 1, 2 or 3. The causal SNPs were chosen uniformly at random among the available SNPs in the region. The causal (non-zero) SNP effects $b_j$ were drawn randomly from the standard normal, then the residual variance $\sigma^2$ was adjusted so that the genotypes at all SNPs in the region explained 0.5% of the variance in $\boldsymbol{y}$. The outcomes $\boldsymbol{y}$ were then simulated as $y_i = x_{i1}b_1 + \cdots + x_{iJ}b_J + \varepsilon_i$, with $\varepsilon_i \sim \mathcal{N}_N(0, \sigma^2)$. We then calculated $z$-scores, $\boldsymbol{z}$, and the in-sample LD matrix $\boldsymbol{R}$ from $\boldsymbol{X}$ and the simulated $\boldsymbol{y}$.

To investigate the impact of misspecification of the LD matrix, we randomly sampled subsets of 500 and 1,000 individuals (not overlapping with the 50,000 samples used to compute the $z$-scores), and computed two "out-of-sample" LD matrices, denoted by $\hat{\boldsymbol{R}}_{500}$ and $\hat{\boldsymbol{R}}_{1000}$, respectively. Because the sample sizes were not large (at most 50,000), it was feasible to compute all in-sample and out-of-sample LD matrices using the function `cor` in `R`.

To assess performance in identifying causal SNPs with larger effects, we repeated the same simulations as described above, except that (a) we used a smaller number of samples, and (b) $\sigma^2$ was adjusted so that the genotypes at all SNPs explained a larger proportion variance in $\boldsymbol{y}$. Specifically, we repeated simulations at target PVE settings of 10% and 30%, and for these settings we used 2,500 samples and 800 samples, respectively, to achieve roughly the same level of power as the simulations with 0.5% and 50,000 samples.

##### Fine-mapping methods

In the simulations, all *SuSiE-RSS* variants were implemented by calling function `susie_rss` from the R package `susieR` (version 0.12.06). Unless otherwise stated, we set the maximum number of non-zero effects to 10 (`L = 10`), we fixed the residual variance $\sigma^2$ to 1 (`estimate_residual_variance = FALSE`, `residual_variance = 1`), we set the maximum number of IBSS-ss iterations to 1,000 (`max_iter = 1000`), and we used refinement (`refine = TRUE`). In all calls to `susie_rss`, the summary data provided were the $z$-scores, LD matrix, and sample size (passed via `susie_rss` arguments `z`, `R` and `n`). We considered the following variants of *SuSiE-RSS*: `susie_rss` with `L`, the maximum number of non-zero effects in the regression model, set to 10 or the true value (1, 2 or 3); with and without the refinement procedure (`refine = TRUE`, `refine = FALSE`); and estimating the residual variance $\sigma^2$ or fixing it to 1 (`estimate_residual_variance = TRUE` or `FALSE`). Note that *SuSiE-RSS* with sufficient data $(\boldsymbol{R}, \hat{\boldsymbol{z}}, N)$ and `estimate_residual_variance = TRUE` (second row of Table 1) gives the same result as *SuSiE* with individual-level data.

We also assessed performance of FINEMAP [6] (version 1.4), CAVIAR [4] (version 2.2) and DAP-G [16, 17] (git commit id 875ba40). All these methods accept summary data $\boldsymbol{z}, \hat{\boldsymbol{R}}$ as input. FINEMAP additionally requires the sample size $n$. (DAP-G accepts either sufficient statistics or summary data, but in our simulations we only evaluated DAP-G with summary data $\boldsymbol{z}, \hat{\boldsymbol{R}}$.) These methods are based on the same multiple linear regression model as *SuSiE-RSS*, differing in the choice of priors, the approach used to compute posterior probabilities, and definition of a credible set.

We ran the FINEMAP program with flags `--sss --n-causal-snps 5`; with these options, FINEMAP used shotgun stochastic search to explore causal configurations, restricting to configurations with at most 5 causal SNPs (this is also the default setting in FINEMAP 1.4). For the "FINEMAP, $L =$ true" results, we instead called FINEMAP with `--sss --n-causal-snps L`, where `L` was the number of causal SNPs used in the simulation (1, 2 or 3). Credible sets were introduced in newer versions of FINEMAP. In FINEMAP version 1.4, a credible set was defined conditioned on the number of causal SNPs, $k$: "For a specific $k$, FINEMAP takes the $k$-SNP causal configuration with highest posterior probability and then asks, for the $l$th SNP in that set, which are the other candidates that could possibly replace that SNP in this causal configuration. The $l$th credible set shows the best candidate SNPs and their posterior probability of being in a $k$-SNP causal configuration that additionally contains $k-1$ SNPs. Note that the $k-1$ SNPs are chosen to have highest posterior probability in their credible set." FINEMAP outputted a set of results for each $k = 1, \ldots, L$. We kept the credible sets from the $k$ with the highest posterior probability.

We ran DAP-G program using the default settings. Note that the default maximum number of causal SNPs in DAP-G (the "maximum model size") is $J$, the total number of SNPs. The DAP-G software outputs "signal clusters" [16], not credible sets. However, we were able to compute credible sets from the DAP-G output following this description from [16]: "For a signal whose local fdr $\leq t$, it is straightforward to construct a $(1-t)$% Bayesian credible set by selecting a minimum subset of SNPs such that their cumulative SNP-level PIPs reach $1-t$." We implemented this calculation in R to generate the DAP-G credible sets.

We ran CAVIAR with flags `-g 0.001 -c L` so that all SNPs had a prior inclusion probability of 1/1000, and the maximum number of causal SNPs was $L$, where $L$ was the value used to simulate the phenotypes (1, 2 or 3). The remaining CAVIAR parameters were kept at their default settings.

To compare CSs generated by *SuSiE-RSS*, FINEMAP and DAP-G (Fig. 4), we first filtered out all CSs with purity less than 0.5 (this is the default setting in `susieR`). Following [1], "purity" is defined as the smallest absolute correlation among all pairs of SNPs within a CS.

##### Computing environment

All simulations were run on Linux machines (Scientific Linux 7.4) with Intel Xeon E5-2680v4 ("Broadwell") processors. *SuSiE-RSS* was run in R 3.6.1 [18] linked to the OpenBLAS 0.2.19 optimized numerical libraries. At most 2 GB of memory was needed to run *SuSiE-RSS* on the simulated data sets, and at most 10 GB was needed to run DAP-G, FINEMAP and CAVIAR. All methods and other computations were run without mutlithreading (one CPU). Runtime statistics for running the methods on summary data with in-sample LD matrices are given in Table 2.

We used the Dynamic Statistical Comparisons system (https://github.com/stephenslab/dsc) to perform the simulations. All code implementing the simulations, and the raw and compiled results generated from our simulations, are available at https://github.com/stephenslab/dsc_susierss, and were deposited on Zenodo [19].

#### References


1. Wang G, Sarkar A, Carbonetto P, Stephens M. A simple new approach to variable selection in regression, with application to genetic fine mapping. Journal of the Royal Statistical Society, Series B. 2020;82(5):1273–1300.

2. Servin B, Stephens M. Imputation-based analysis of association studies: candidate regions and quantitative traits. PLoS Genetics. 2007;3(7):e114.

3. Chipman H, George EI, McCulloch RE. The practical implementation of Bayesian model selection. In: Model Selection. vol. 38 of IMS Lecture Notes. Institute of Mathematical Statistics; 2001. p. 65–116.

4. Hormozdiari F, Kostem E, Kang EY, Pasaniuc B, Eskin E. Identifying causal variants at loci with multiple signals of association. Genetics. 2014;198(2):497–508.

5. Kichaev G, Yang WY, Lindstrom S, Hormozdiari F, Eskin E, Price AL, et al. Integrating functional data to prioritize causal variants in statistical fine-mapping studies. PLoS Genetics. 2014;10(10):e1004722.

6. Benner C, Spencer CCA, Havulinna AS, Salomaa V, Ripatti S, Pirinen M. FINEMAP: efficient variable selection using summary data from genome-wide association studies. Bioinformatics. 2016;32(10):1493–1501.

7. LaPierre N, Taraszka K, Huang H, He R, Hormozdiari F, Eskin E. Identifying causal variants by fine mapping across multiple studies. bioRxiv. 2020;doi:10.1101/2020.01.15.908517.

8. Albert A. Regression and the Moore-Penrose pseudoinverse. New York, NY: Academic Press; 1972.

9. Trefethen LN, Bau D. Numerical linear algebra. Philadelphia, PA: SIAM; 1997.

10. Maller JB, McVean G, Byrnes J, Vukcevic D, Palin K, Su Z, et al. Bayesian refinement of association signals for 14 loci in 3 common diseases. Nature Genetics. 2012;44(12):1294–1301.

11. Brent RP. Algorithms for minimization without derivatives. Mineola, NY: Dover; 2002.

12. Kim Y, Carbonetto P, Stephens M, Anitescu M. A fast algorithm for maximum likelihood estimation of mixture proportions using sequential quadratic programming. Journal of Computational and Graphical Statistics. 2020;29(2):261–273.

13. Sudlow C, Gallacher J, Allen N, Beral V, Burton P, Danesh J, et al. UK Biobank: an open access resource for identifying the causes of a wide range of complex diseases of middle and old age. PLoS Medicine. 2015;12(3):e1001779.

14. Bycroft C, Freeman C, Petkova D, Band G, Elliott LT, Sharp K, et al. The UK Biobank resource with deep phenotyping and genomic data. Nature. 2018;562(7726):203–209.

15. Canela-Xandri O, Rawlik K, Tenesa A. An atlas of genetic associations in UK Biobank. Nature Genetics. 2018;50(11):1593–1599.

16. Lee Y, Luca F, Pique-Regi R, Wen X. Bayesian multi-SNP genetic association analysis: control of FDR and use of summary statistics. bioRxiv. 2018;doi:10.1101/316471.

17. Wen X, Lee Y, Luca F, Pique-Regi R. Efficient integrative multi-SNP association analysis via deterministic approximation of posteriors. American Journal of Human Genetics. 2016;98(6):1114–1129.

18. R: a language and environment for statistical computing; 2018. Available from: https://www.R-project.org.

19. Zou Y, Carbonetto P, Wang G, Stephens M. stephenslab/dsc_susierss: Release of dsc_susierss repository accompanying journal resubmission; 2022. Available from: https://doi.org/10.5281/zenodo.5611713.
