## Supplementary figures and images for "Fine-mapping from summary data with the “Sum of Single Effects” model"

### Supporting Figure S1

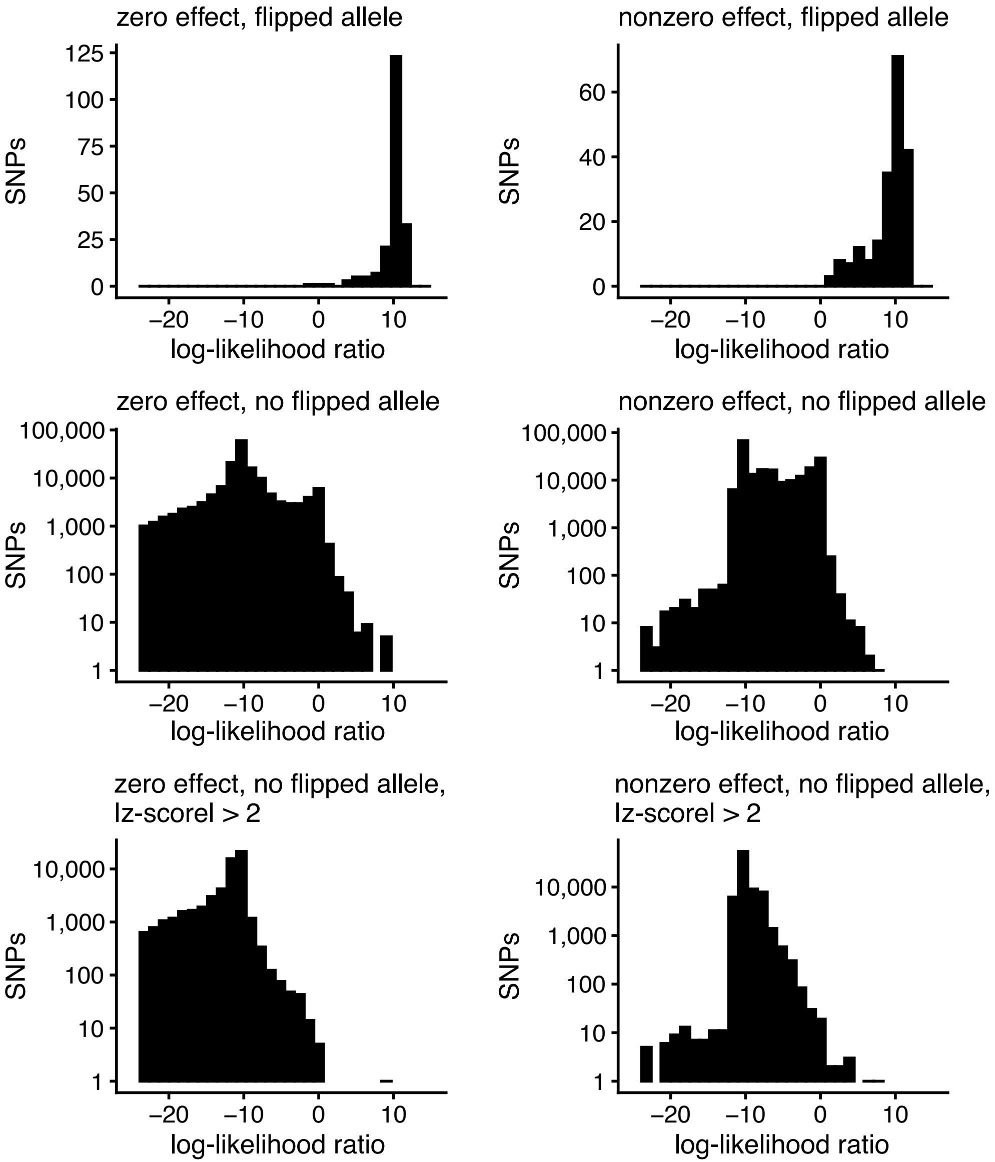
